# Molecular interactions underlying selective partitioning of clients into MED1 and FUS condensates

**DOI:** 10.64898/2025.12.23.696191

**Authors:** Ikki Yasuda, Eiji Yamamoto, Kenji Yasuoka, Kresten Lindorff-Larsen

## Abstract

Biomolecular condensates selectively recruit and exclude proteins, and this partitioning is sometimes dictated by their intrinsically disordered regions. Factors that determine partitioning include client–scaffold interactions and stabilisation of the scaffold network within the condensate, yet a molecular and predictive understanding has not been fully achieved. Here, we used coarse-grained molecular dynamics simulations to elucidate molecular interactions between two types of nuclear proteins as scaffolds: the charge-rich disordered region of Mediator 1 (MED1) and the aromatic-rich disordered region of Fused in sarcoma (FUS), in mixtures with client intrinsically disordered regions from six different proteins. We show distinct partitioning into the MED1 and FUS condensates, while enrichment of RNA in the dilute phase modulated partitioning in FUS condensates. Favourable client–scaffold interaction energies within condensates were associated with client partitioning, while client–scaffold interactions competed with scaffold self-interactions. Analysis of interaction energies of individual residues revealed that the interactions were localised to specific sequence regions: charged blocks for interactions with MED1, and dispersed aromatic residues for interactions with FUS. Based on these molecular insights, we developed a sequence-based predictor of partitioning trends and applied it to a set of 243 sequences. Our prediction approach for partitioning can be extended to other biomolecules, offering a framework to analyse their partitioning in cellular environments.

## 1 Introduction

Biomolecular condensates, biomolecular assemblies formed via multivalent interactions among biomolecules, are thought to play important roles in the regulation of various cellular activities such as transcription, signal transduction, and biomolecular degradation.^1–3^ A central function of biomolecular condensates is to recruit biomolecules and small molecules through their unique physicochemical environments, thereby enabling the spatial organization of biomolecules without membranes.^4–8^ Partitioning of biomolecules can directly control biochemical processes,^9–11^ potentially relating to the development of diseases.^12,13^ Furthermore, partitioning also enables demixing of biomolecular condensates within cells, ^14^ leading to coexistence of diverse types of membraneless organelles.^15,16^ However, it is still difficult to predict which molecules will selectively partition into a given condensate.

The selective partitioning of biomolecules is primarily governed by molecular interactions within condensates.^17–21^ Thus, understanding the interactions between scaffold molecules, which form the condensate framework, and client molecules, which partition into them, is essential to explain selectivity.^22–24^ Early studies in partitioning focused on model systems composed of repeats of folded domains and their cognate binding partners.^23,25^ In these systems, partitioning was governed by availability of interaction sites, and clients with stronger affinity for the scaffold sites effectively outcompeted and displaced weaker binders.^23^ In contrast, individual interactions via amino acid residues in intrinsically disordered regions (IDRs) are typically weak and non-specific, yet via multivalency, they can drive biomolecular condensation.^26^ IDR condensates can also recruit clients, including specific proteins with characteristic sequence composition or charge patterning,^9,20,27,28^ as well as disordered nucleic acids.^12,29^ The resulting selectivity is governed by a complex interplay of factors, such as scaffold–client interactions, partial disruptions of scaffold networks,^29^ diffusive properties,^30,31^ and the balance of enthalpy and entropy.^32,33^ Due to the unstructured nature of IDRs, the molecular recognition mechanism of IDR condensates can differ from structurebased recognition mediated by folded domains.^34,35^

Recently, De La Cruz *et al.* reported the differential partitioning of transcription factor IDRs into condensate formed by either a charged-residue-rich IDR from Mediator 1 (MED1), or an aromatic-residue-rich IDR from Fused in sarcoma (FUS), in *in-vitro* and *in-vivo* measurements.^36^ The MED1 clients are generally abundant in negatively charged residues, while the FUS clients are rich in aromatic residues, but some IDRs do not partition either of them, while others partition into both.^36^ Therefore, the electrostatic and hydrophobic interactions are one of the primary factors in the selective partition of IDRs, which nevertheless does not explain all of the partitioning trends.^28^ Despite progress in discovering the molecular basis for the rules underlying partitioning and mixing,^20,28,37–39^ details of the molecular grammars of heterotypic interactions and the balance between homotypic and heterotypic interactions among scaffolds and clients, remain unclear.

Coarse-grained molecular dynamics (MD) simulations at one-bead-per-residue resolution have been used to provide insights on the molecular interactions within biomolecular condensate since these models can be used to describe large-scale systems, with maintaining some level of the chemistry of biomolecules.^40^ Generally, these one-bead-per-residues models contain both non-ionic ‘stickiness’-based attractive interactions and electrostatic interactions, which partially capture the driving forces of condensation such as electrostatic, *π* interactions and hydrophobic interactions, combined with implicit solvation.^41–43^ Previously, we have optimized sequence-dependent amino acid stickiness parameters of the hydropathy scale model to reproduce experimental values of single-chain properties of various IDRs using Bayesian regularization approach,^44^ and shown that the optimized model with useful accuracy predicted condensation properties.^45–47^ We also developed a compatible coarse-grained model for disordered RNA, which captures some aspects of both polymer properties and phase separation properties with IDRs.^48^ Therefore, these models enable simulations of mixed protein–RNA condensates as well as protein condensates.

Our work aims to elucidate the molecular mechanisms underlying the selective partitioning of nuclear protein IDRs into MED1 and FUS IDR condensates. Using coarse-grained MD simulations of mixtures of IDR condensate and client IDRs, we demonstrate that our simulations capture several key aspects of the selective partitioning observed in previous experiments.^36^ Through detailed analyses of the molecular interactions, we find that clients interact with scaffold proteins and RNA in a sequence-dependent manner, identifying the specific sequence regions involved in these interactions. Finally, leveraging these insights on the localised interactions, we develop a sequence-based predictor for the selective partitioning behaviour into MED1 and FUS condensates.

## 2 Results

### 2.1 Client IDR partitioning into MED1 and FUS condensates

In this study, we focused on two scaffold proteins, the C-terminal IDR of MED1 (UniProt ID Q15648; residues 948–1573) and the N-terminal IDR of FUS (UniProt ID P35637; residues 2–214). Distinct client recruitment by these two IDR condensates has been observed in both *in-vitro* and *in-vivo* experiments.^36^

MED1 is a transcriptional coactivator and forms transcriptional condensates together with nucleic acids, transcription factors and RNA polymerase II.^49,50^ The long C-terminal IDR of MED1 is highly positively charged and therefore does not undergo phase separation on its own, but can do so in the presence of crowding agents or nucleic acids.^51,52^

In contrast, the N-terminal IDR of FUS, which is enriched in tyrosine residues, shows robust condensate formation under physiological conditions both *in vitro* and *in vivo*.^36,53^ Contrary to full-length FUS, whose phase separation is biphasically affected by RNA,^54^ its N-terminal condensates exclude disordered RNA.^53^

As client proteins, we selected six IDRs (from SPT6, CTR9, CDK11B, EWSR1, KHDR1, and p300) based on the study by De La Cruz *et al.*^36^ Each client type was included individually in separate simulations, such that each simulation contained only one type of client IDR. These client IDRs span a broad sequence space in terms of charge and aromatic content (Fig. 1a) and exhibit differential partitioning into MED1 and FUS condensates.^36^

**Figure 1:**
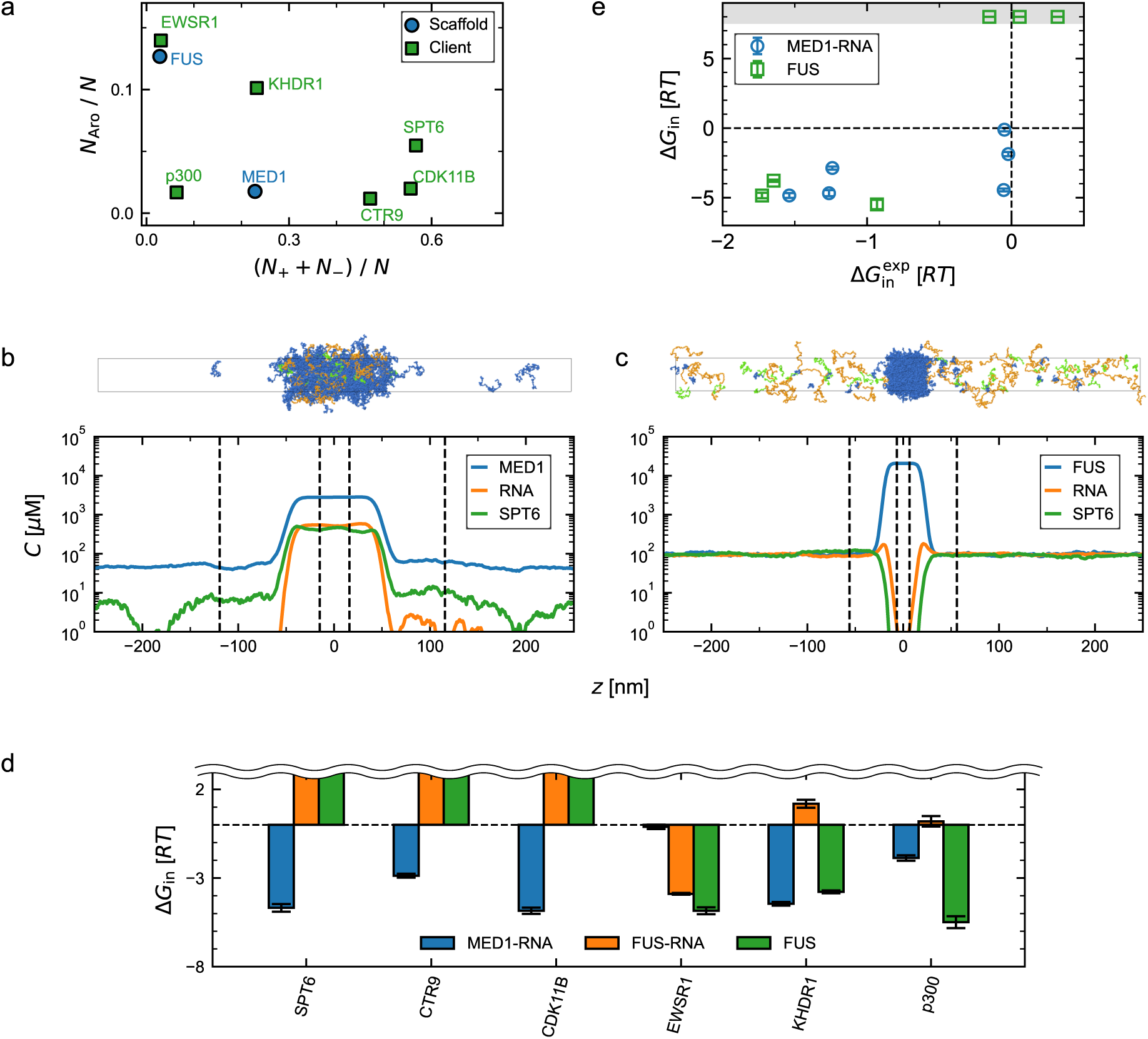
Selective partitioning of clients into Mediator 1 (MED1) and Fused in Sarcoma (FUS) condensates. (a) Mapping of host and client IDRs based on charged and aromatic residue fractions. *N*, *N*_+_, *N_−_*, and *N*_Aro_ denote the sequence length, the number of positively charged residues, the number of negatively charged residues, and the number of aromatic residues, respectively. (b, c) Representative snapshots and corresponding density profiles from MD simulations of (b) MED1 condensate and (c) FUS condensate, both using SPT6 as an example client protein. Density profiles for other systems are shown in Fig. S1. (d) Transfer free energy, Δ*G*_in_, of clients into three types of condensates: MED1–RNA condensate (blue), FUS condensate with RNA in the dilute phase (orange), and FUS condensate without RNA (green). Since the Δ*G*_in_ values of highly excluded clients could not be determined with confidence, their precise values are not shown. (e) Comparison between the Δ*G*_in_ and the experimental values derived from *in-vivo* measurements of the partition coefficient.

Our simulations contained scaffold proteins, unstructured single-stranded RNAs, and clients, with the aim of evaluating the extent to which clients partition into scaffold-comprising condensates. Both scaffold proteins (MED1 and FUS) formed condensates in our simulations using the slab geometry, while only MED1 localised together with RNA (Fig. 1b).^48^ The client protein SPT6 partitioned into the MED1 condensate (Fig. 1b), but was excluded from the FUS condensate (Fig. 1c). Similar simulations were performed for all combinations of client and scaffold proteins (density profiles for other systems are shown in Fig. S1).

As a quantitative measurement of client partitioning, client concentrations in the dense, *c*_den_, and dilute phases, *c*_dil_, were obtained from the density profiles, and the resulting concentration ratio was converted into a transfer energy, Δ*G*_in_ = −*RT* ln(*c*_den_*/c*_dil_) with temper-ature *T* and the gas constant *R*. We note that our simulations represent a single point on a multi-dimensional phase diagram, and thus that the Δ*G*_in_ values only quantify the partitioning under the specific conditions and concentrations of protein and RNA. In particular, we note that we use lower concentrations of the ‘client’ so that the simulations mostly represent partitioning into a scaffold-rich condensate environment. The MED1 condensate recruited all six client chains, with particularly strong partitioning observed for charge-rich sequences, whereas non-charge-rich clients such as p300 and EWSR1 showed much weaker enrichment (Fig. 1d). In contrast, the FUS condensate (with RNA present in the dilute phase) strongly recruited EWSR1, slightly excluded p300 and KHDR1, and substantially excluded SPT6, CTR9, and CDK11B (Fig. 1d). Notably, the apparent exclusion of p300 and KHDR1 from the FUS condensate was attributed to their surface localisation, as revealed by their density profiles (Fig. S1), which was not captured in our transfer-energy calculation. De La Cruz *et al.*^36^ identified SPT6, CTR9, and CDK11B as MED1-recruiting clients, while EWSR1, KHDR1, and p300 were classified as FUS-recruiting. Our simulations were thus in qualitative agreement with the MED1-recruitment trend, while some FUS-recruiting clients were apparently excluded from the condensate centre.

We reasoned that the surface enrichment of FUS clients originated from the interaction of the client with RNA present in the dilute phase, and therefore performed additional simulations using RNA-free FUS systems. In the RNA-free FUS systems, FUS condensates recruited EWSR1, p300 and KHDR1, while excluding other clients (Fig. 1d), consistent with the experimental recruitment trend. We speculate that, in the experiments, the dilute phase occupies a large volume fraction, resulting in low RNA concentrations outside of the condensates. Based on this hypothesis, we used the RNA-free FUS condensate as FUS systems in the following analyses, while the MED1 systems contained RNA.

The partitioning energies obtained from our simulations correlated relatively well with experimental partitioning energies derived from *in-vivo* measurements of the partition coefficient^36^ (Fig. 1e; the Spearman rank-order correlation coefficient=0.71). However, the partitioning of SPT6, CTR9, and CDK11B into FUS condensates was not observed during our simulation. Therefore, the dense-phase concentrations of these clients could not be determined quantitatively. For this reason, we assigned a high transfer energy of 8*RT* to these clients in Fig. 1d, but these values should not be regarded as reliable measurements. Again, we note that in the *in-vivo* situation, the condensate composition may differ from that in our simulations, hampering more quantitative comparisons.

### 2.2 Competition between scaffold–scaffold and scaffold–client interactions

Since the partitioning of clients is driven by favourable scaffold–client interactions, we evaluated the client–scaffold interactions within the condensate by calculating the interaction energies between a central chain of the client and the surrounding scaffolds. Here, the central client chain refers to the chain located closest to the centre of the condensate at each time frame, representing the molecular environment inside the condensate (Fig. 2a; see Methods for details). Scaffold–client interactions were defined as intermolecular interactions between the central client chain and the surrounding macromolecules, which were either MED1 or RNA in the MED1 system, and FUS in the FUS system. Calculations were also performed for the homotypic scaffold interactions between the central scaffold chain and the surrounding scaffolds. The scaffold–client interaction energy correlated with the partitioning energy (Fig. 2b), consistent with the previously reported relationship between energetic gain from homotypic interactions and saturation concentration.^55^

**Figure 2:**
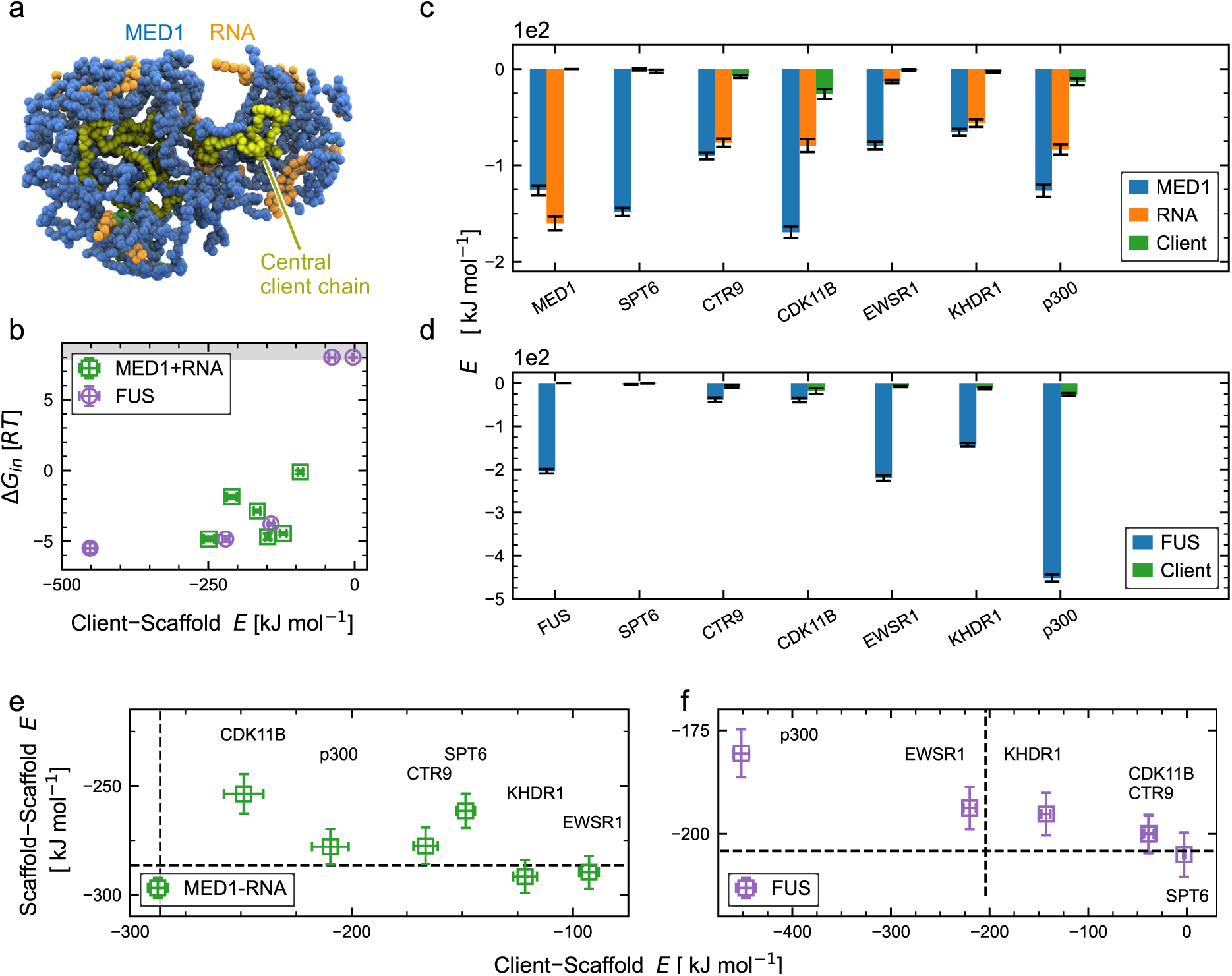
Competitive interaction between scaffold and client chains. (a) Representative snapshot of the central client chain and the surrounding molecules within 4 nm in the MED1–RNA–SPT6 system. The central client chain is shown in yellow, the scaffold chain in blue, RNA in orange, and surrounding client chains in green. (b) Partition energy from MD simulations, Δ*G*_in_, plotted against Client–Scaffold energy *E*, which we calculated as the sum of client–MED1 and client–RNA interactions, or the client–FUS interactions. (c, d) Intermolecular interaction energy of the central client chain, *E*, computed for client–scaffold protein, client–RNA, and intermolecular client–client interactions. Condensate of MED1 in (c) and FUS in (d). The x-labels ‘MED1’ and ‘FUS’ indicate scaffold–scaffold interactions from client-free systems, whereas the other labels indicate clients. (e, f) Interaction energy of the central client chain, plotted against that of the central scaffold protein chain. Scaffold– scaffold protein interaction energy, Scaffold–Scaffold *E*, includes MED1–MED1 and MED1– RNA in (e), and FUS–FUS interactions in (f). Dashed black lines represent the strength of scaffold–scaffold interactions obtained from client-free condensate. Error bars in panels (b–f) represent standard error of mean estimated using the time-blocking approach.

Given that client interactions within the MED1–RNA condensate involve both MED1 and RNA, decomposing the interaction energy by macromolecule type (MED1, RNA, and the surrounding client molecules) helps to clarify the individual contributions. Among the highly partitioning clients, SPT6 and CDK11B exhibited substantially stronger interactions with MED1 than with RNA, whereas CTR9, KHDR1, and p300 showed comparable levels of interaction with both MED1 and RNA (Fig. 2c). These results suggest that direct interactions with MED1 predominantly drive partitioning for the former group, while RNA also plays a non-negligible role for the latter. As a control, we performed additional simulations of client-free binary systems and found that MED1–RNA interactions are stronger than MED1–MED1 interactions (Fig. 2c), indicating that the presence of RNA enhanced the overall interaction energy and promoted condensate formation.^48,52^ A similar decomposition of interaction energies in binary mixtures of FUS and individual clients indicates that partitioning clients (EWSR1, KHDR1, and p300) exhibit strong interactions with FUS (Fig. 2d). Furthermore, in our simulations, a low client concentration was maintained, resulting in minimal intermolecular interactions among client molecules across all simulations.

We reasoned that heterotypic client–scaffold interactions stronger than homotypic scaffold– scaffold interactions would enable client partitioning. Scaffold–scaffold interactions are defined as the intermolecular interaction energy between a central scaffold chain and the surrounding scaffold chains, calculated using the central-chain protocol described above. The client–scaffold interaction energy was negatively correlated with the scaffold–scaffold interaction energy (Figs. 2e and f), indicating that client interactions partially replaced homotypic scaffold interactions.

Contrary to our expectation, in MED1 condensate, scaffold–scaffold interactions were substantially stronger than client–scaffold interactions for all tested clients (Fig. 2e). We interpret that the MED1 condensates were not tightly packed, allowing client partitioning without substantially disrupting scaffold interactions. To support this interpretation, we calculated the particle number densities of scaffold molecules in the dense phase. The average density over all MED1 systems was 1.2 particles*/*nm^3^, which was less than half that of the FUS condensates of 2.5 particles*/*nm^3^. In the tightly packed FUS condensate, only clients with sufficiently strong interactions relative to scaffold–scaffold interactions appear able to partition into the condensate. FUS-client interactions exceeded scaffold–scaffold interactions for p300 and EWSR1 (Fig. 2f), whereas KHDR1 exhibited weaker interactions yet partitioned moderately, likely due to its shorter sequence length.

These results suggest that client partitioning arises when heterotypic client–scaffold interactions exceed homotypic scaffold–scaffold interactions,^29,33^ or when the client interacts weakly but causes only minimal disruption of the scaffold network. The latter case applies to all clients in MED1 condensates and to KHDR1 in FUS condensates.

### 2.3 Client–scaffold interactions localize at specific sequence regions

The sequence patterning of client IDRs is heterogeneous, exhibiting features such as charge block formation and dispersed aromatic residues. Here, we identify the specific regions within the sequence that contribute to client–scaffold interactions. Instead of commonly used contact maps, we employed per-residue interaction energy profiles to maintain consistency with our preceding interaction energy analysis (Fig. 2c and d).

We identified the central client chain and calculated the per-residue interaction energies by summing the interactions of each client residue over all surrounding scaffold molecules. SPT6, which contains both positively and negatively charged blocks, strongly interacted with MED1 (interaction energy *>* 1 kJ*/*mol) via its positively charged regions (Fig. 3a), supporting the importance of such blocks in selective partitioning.^9^ A previous study reported similar per-residue interaction energies of approximately 1 kJ mol*^−^*^1^ bead*^−^*^1^ within condensates using a similar coarse-grained model.^55^ CTR9, which also contains charge blocks, interacted with RNA through its positively charged region spanning residues 900–1000, whereas the C-terminal region exhibited only weak interactions with MED1. In the MED1–RNA condensate, KHDR1 interacted with RNA at the terminal regions enriched in sparsely distributed positive charges, while its central region (residues 360–420), enriched in aromatic and acidic residues, showed weak interactions with MED1. Within the FUS condensate, EWSR1 exhibited strong interactions with FUS through its aromatic residues distributed across its sequence (Fig. 3d). Sequence patterning influenced the localisation of interactions, which was characteristic of each individual sequence (residue interaction profiles for the other MED1 and FUS systems are shown in Figs. S2 and S3, respectively).

**Figure 3:**
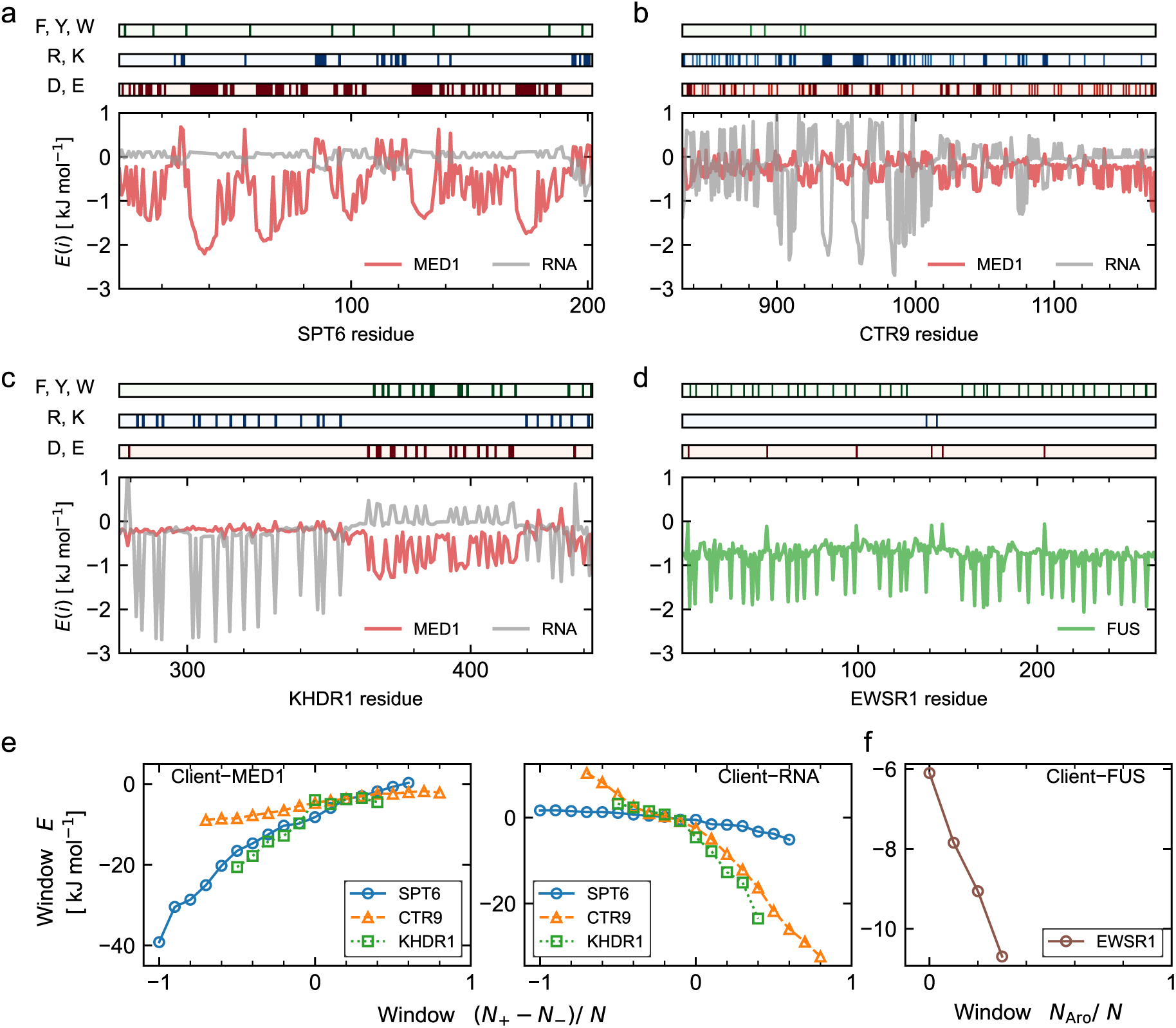
Analysis of client–scaffold interactions decomposed by client residues. (a–d) Interaction energy of residue *i*, *E*(*i*). Aligned sequence plots highlight aromatic (F, Y, W), positively charged (R, K), and negatively charged (D, E) residues. Scaffold protein–client and RNA–client interactions are shown for MED1–RNA–client system in (a), MED1–RNA– CTR9 system in (b), MED1–RNA–KHDR1 in (c), and FUS–EWSR1 system in (d). Residue interactions profiles for the other MED1 and FUS systems are shown in Figs. S2 and S3, respectively. (e, f) The sum of residue interaction energies over ten-residue windows, window *E*, is computed. For each value of the net charge per residue, the mean of window *E* across all windows is then calculated. Client–MED1 or client–RNA interactions for clients (SPT6, CTR9, KHDR1) are shown in (e), and client–FUS interaction for EWSR1 is shown in (f).

We then evaluated how residue clustering influences interaction energies. We obtained ten-residue windows along the sequence from the per-residue interaction energy profiles, summed *E*(*i*) within each window, and compared these with the net charge per residue (NCPR) for MED1 or the fraction of aromatic residues for FUS systems. The average over multiple windows was used to represent the relationship between the degree of residue clustering and interaction energy.

In client–MED1 interactions, we observed that windows with negative NCPR tend to exhibit substantially stronger interactions, whereas windows with positive NCPR showed near-zero interaction energy (Fig. 3e). Client–RNA interactions showed a similar trend but with the opposite charge preference (Fig. 3e). In FUS–EWSR1 interactions, the fraction of aromatic residues per window was relatively low (*<* 0.4), and, within this range, the interaction energy linearly increased (Fig. 3f). These results indicate that charge patterning causes significantly strong local interactions.

### 2.4 Sequence-based client partitioning predictor accounting for interaction localisation

Based on the observed relationship between composition, sequence patterning and localised interactions, we subsequently developed a simple sequence-based predictor for client partitioning. Previous studies have proposed approaches to predict intermolecular interactions based on residue potential functions of coarse-grained molecular models,^28,56,57^ and charge patterning.^39^ Building on these ideas, we propose a sequence-block interaction framework, in which sequences are segmented based on sticker patterning, and the interactions between the resulting blocks are quantified.

In the sequence-block interaction framework, we first defined blocks of sticker residues based on the CALVADOS 2 potential (Fig. 4a), categorizing residues into three types (basic, acidic, and aromatic). Blocks were formed by grouping neighbouring residues of the same type, thus representing the SPT6 sequence as a series of acidic blocks (Fig. 4a). Subsequently, we defined the block–block potential, *U_ij_*(*r*), using the pair-wise potential, *u_kl_*(*r*),

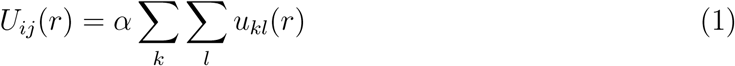

where *i* and *j* is the indices of blocks, and *k* and *l* are the indices of residues of blocks *i* and *j*, respectively. The potential, *u_kl_*(*r*), is the sum of Ashbaugh–Hatch (AH) and Debye-Hückel (DH) potential of residue *k* and *l* at distance *r*. The scaling parameter *α* was set to 3 to emphasize strong interactions. We scanned the parameters related to sequence blocking and the energy scaling factor, *α*, so that *B*^tot^ captures the distinct partitioning behaviour between MED1 and FUS. The interaction between the block *i* and *j* is quantified using the second virial coefficient,

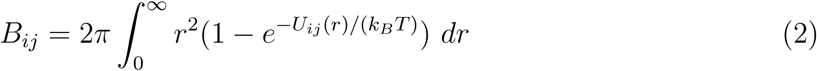

**Figure 4:**
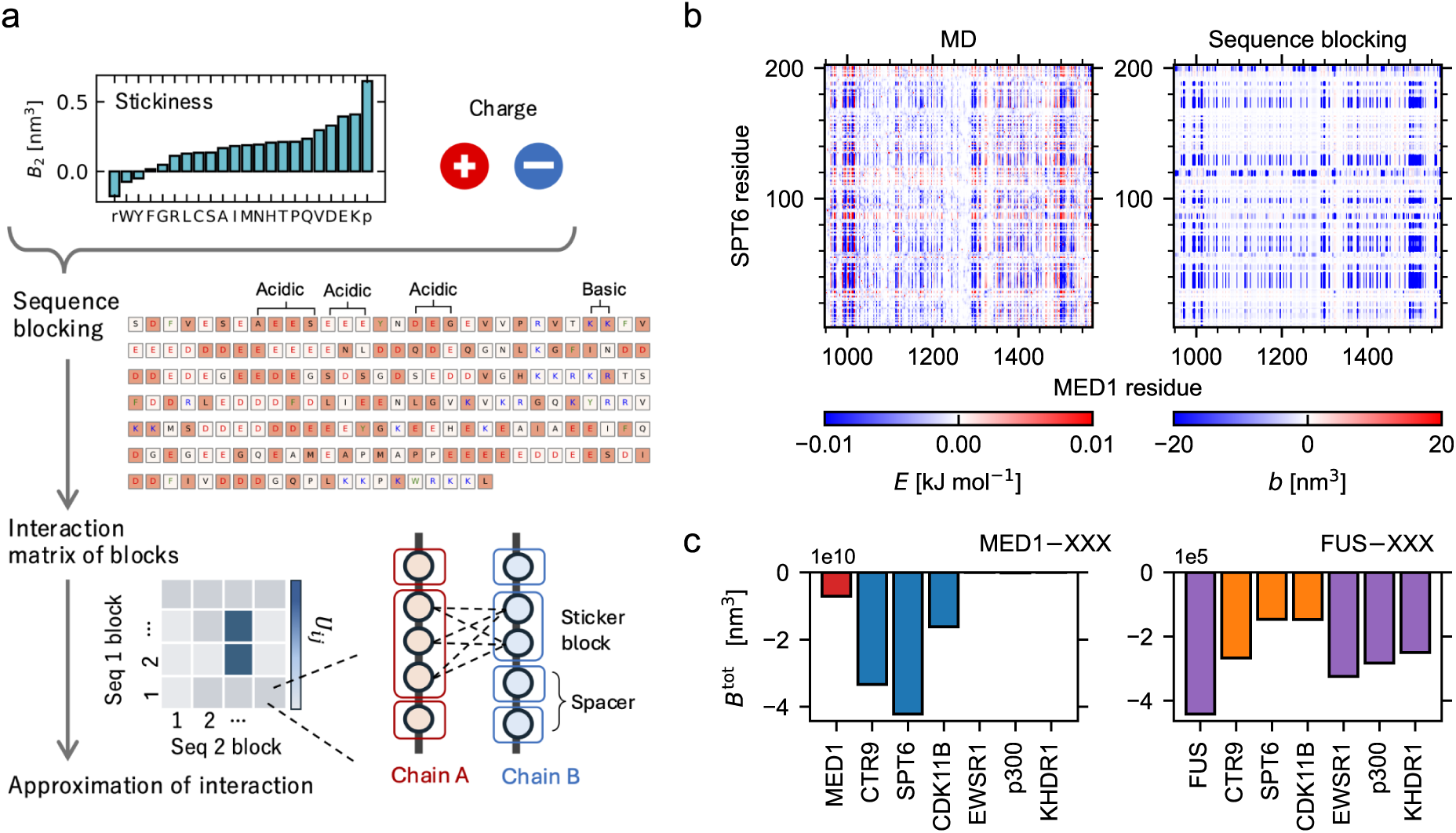
Sequence-base prediction of intermolecular interactions. (a) Overview of the computational scheme. (b) Comparison of residue–residue interactions between MED1 and SPT6, based on the slab MD simulation in the presence of RNA and the sequence-blocking approach applied to the MED1–SPT6 pair. (c) Prediction of the intermolecular interaction parameter from the blocking approach, block *B*. Calculations were performed by fixing the protein (MED1 and FUS), while varying the client. For the interactions of MED1, red/blue bars show unfavourable/favourable interactions with MED1, respectively. Purple/orange bars show favourable/unfavourable interactions with FUS, respectively.

The second virial coefficient act as a nonlinear transformation of the block–block potential. Therefore, this computation can capture the pronounced interactions that occur at sticker clusters. Finally, the sequence–sequence interaction is computed as the sum of the block interactions,

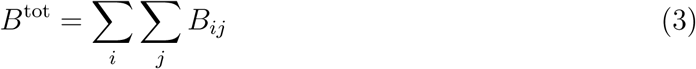

We stress that the purpose of *B*^tot^ is to capture the partitioning behaviour of client chains into scaffold IDRs condensates, rather than to compute the exact second virial coefficient.

To test whether our predictor captures scaffold–client interactions, we first mapped the *B_ij_* values onto pairwise-residue interactions, assigning each residue pair an average interaction value, *b* = *B_ij_/n_i_/n_j_*, where *n_i_* and *n_j_* are the number of residues in each block. We then compared this predicted map with the contact energy map obtained from our MD simulations (Fig. 4b). Our predictor successfully captured the attractive interaction sites in MED1 around residues 970, 995, 1110, 1300, 1500–1520, as well as the overall interaction pattern in SPT6. However, it failed to capture some of the repulsive interactions in SPT6 near residues 90 and 110, likely because these residues contained positively charged blocks that preferentially interacted with RNA within the MED1–RNA condensate (Fig. 3a).

Subsequently, *B*^tot^ values between clients and either MED1 or FUS, as well as the self-interactions of the scaffolds, were compared. MED1 clients (SPT6, CTR9, and CDK11B) showed low *B*^tot^ values, indicating attractive interactions, compared to non-MED1 partitioning proteins and self-MED1 interactions (Fig. 4c). Similarly, FUS clients (EWSR1, p300, and KHDR1), as well as FUS self-interactions, showed low *B*^tot^ values relative to two non-partitioning clients (SPT6, and CDK11B). These results demonstrate that our *B*^tot^ captures some aspects of the relative partitioning behaviour of the six clients.

Having tuned model parameters and validated our predictor on a small client dataset, we then applied it to the MED1/FUS partitioning protein sequence dataset comprising 243 sequences provided by De La Cruz *et al.*^36^ This dataset categorizes sequences from nuclear extracts into three groups based on *in-vivo* and *in-vitro* partitioning behaviour, namely (1) those that partition into both MED1 and FUS, (2) those that partition into MED1 only, and (3) those that partition into FUS only under the conditions of the experiments.

Our predictor captured the distinct trends among the three categories (Fig. 5) and, when used as a classifier of partitioning, performed at least as well as a previously proposed model^56^ as measured by AUC (Fig. S4). We note that the FINCHES method was not developed to predict mixing in systems with RNA, nor to compare proteins with substantial differences in lengths. Nevertheless, we evaluated these predictors to classify four classes: common clients, MED1 clients, FUS clients, and non-clients. The Matthews correlation coefficient (MCC) of the predictor was 0.18 using the thresholds shown in Fig. 5, again performing comparably to the FINCHES method (Tab. S1). We speculate that interactions between the folded domains and the interplay of multiple IDRs occur in the experiments, and quantitative comparisons are also hampered by differences in compositions in the different condensate environments. These complexities make it challenging to accurately predict client partitioning solely from their IDRs, which, in part, explains the moderate performance of the models.

**Figure 5:**
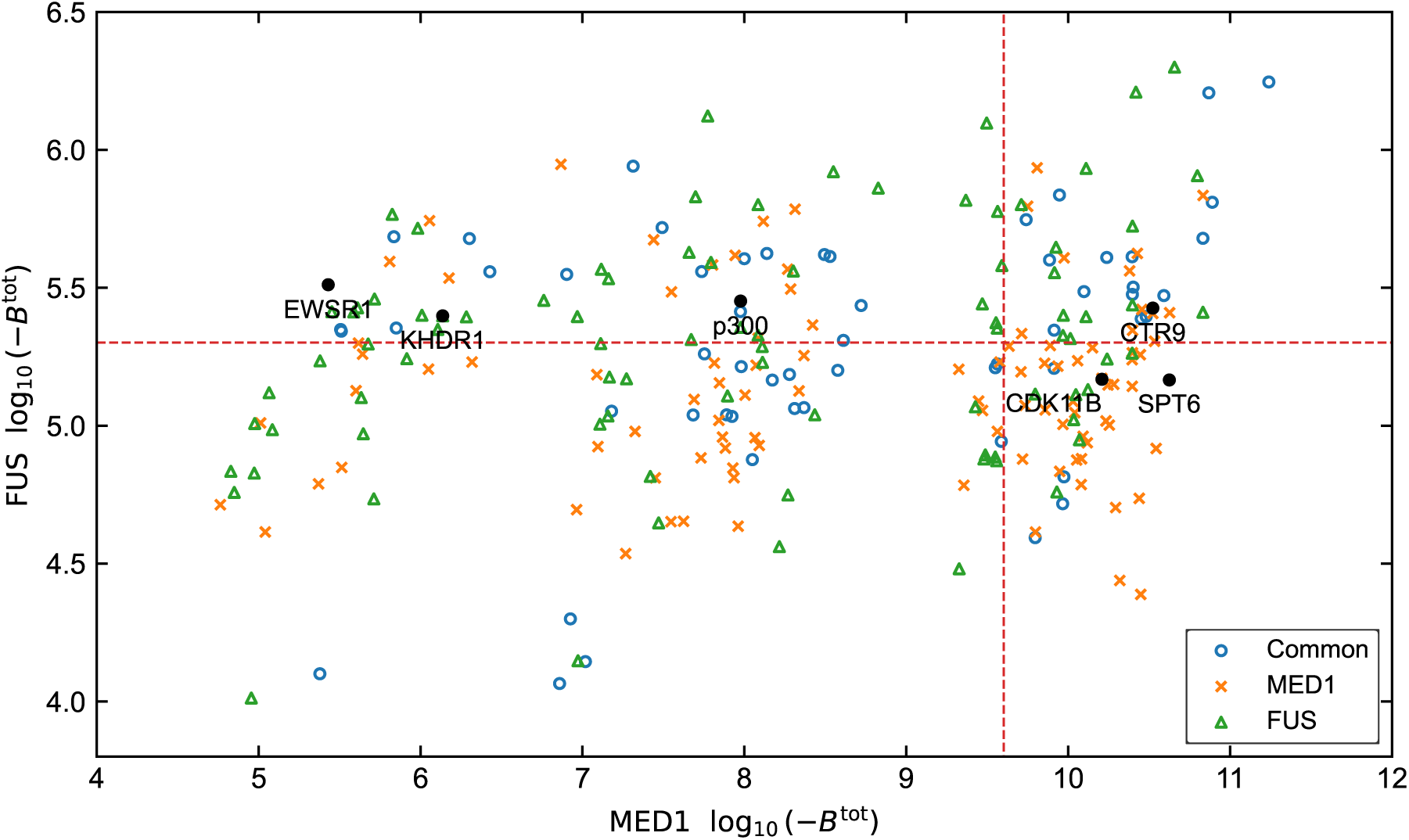
Application of the sequence-based predictor to a large set of IDR sequences. Protein sequences are plotted in the space of the predicted partitioning scores with MED1 (*x*-axis) and FUS (*y*-axis), using a logarithmic scale for visualization, log10(−*B*^tot^). Sequences are grouped into those that partition into both MED1 and FUS (Common, 56 sequences), those partition into MED1 but not FUS (MED1, 100 sequences), and those partition into FUS but not MED1 (FUS, 87 sequences). Red dotted lines indicate thresholds for client classification, which were selected from those evaluated in the ROC curve analysis (Fig. S4).

## 3 Discussion

To elucidate the molecular mechanism underlying the selective partitioning of disordered proteins into biomolecular condensates, we performed coarse-grained MD simulations of MED1 and FUS IDR condensates with client IDRs. We analysed the relationship between the partitioning free energy and the molecular interactions of the clients and scaffolds, and identified residues of the clients that contributed to these interactions. Building on these molecular insights, we developed a sequence-based predictor for client partitioning into the two condensates.

Our results indicate a correlation between the partitioning free energies of the clients and the client–scaffold interaction energies. From Fig. 2b, we can see that for both FUS and MED1 this correlation can be explained by a single curve. This indicates that our client–scaffold interaction energy primarily drives the selective partitioning in the MED1 and FUS condensates. However, we note that the interaction energies, computed from the potential of the coarse-grained model, include both the enthalpic contributions from amino acids and nucleic acids and the effective free energy due to solvents and ions.^58^ For this reason, our interaction energies do not directly represent the actual enthalpic contributions.^32^ Despite this limitation, these interaction energies help rationalize and predict the partitioning behaviour for MED1 and FUS condensates observed in experiments.

We have demonstrated the competition between the scaffold–scaffold interactions and the scaffold–client interactions (Figs. 3c and d). Client partitioning modulates the scaffold networks within the condensates.^29,33^ In our FUS condensates, partitioning clients such as EWSR1 and p300 exhibited scaffold–client interaction energies that exceeded the scaffold– scaffold interactions. However, this competition does not necessarily imply that the partitioning requires scaffold–client interactions to exceed scaffold–scaffold interactions. In loosely packed condensates, clients can be recruited without strongly disrupting scaffold networks within condensates, as observed in our MED1 condensates and our previous simulations of condensates of the disordered region of TDP-43.^59^ Therefore, the condensate density is an important quantity for predicting the partitioning ability of IDR condensates, extending our previous proteomic studies of IDRs that revealed single-chain properties ^60^ and condensate properties such as transfer energies and dilute phase concentrations.^61^

Finally, our results suggest that the composition of the dilute phase also modulates the client partitioning. Our simulations of FUS condensates demonstrated the surface preferences of the clients when RNAs were present in the dilute phase (Fig. 1d). We speculate the dilute phases within the cells may have a similar modulating effect, and partitioning trends vary among cell types and depend on local RNA expression levels and post-translational modifications^62–64^

Our sequence-based prediction model accounts for sequence clustering to emphasize these interactions. Prior studies have introduced energy scales that incorporate local electrostatic properties and hydrophobicity,^56^ as well as models that rely on patterns of adjacent residues^57^ or more general sets of features.^28^ These approaches can parametrise interaction strengths and are particularly effective in capturing mutational effects. However, it remains unclear how well they capture interactions across diverse sequences of varying length and composition. Here, as an extension of the fixed-window approach by Adachi and Kawaguchi,^57^ we proposed a flexible-window-size approach that explicitly accounts for the size and number of analogous residue clusters. As a result, across proteins spanning a wide range of lengths, we extract parameters that quantify partitioning. Our sequence-based predictor is scalable to larger sequence datasets and may be a useful mechanistic supplement for predicting protein subcellular localisation^65^ that complementing other recently described approaches.^66–68^ Our framework is currently limited by (1) the use of a metric that is not fully quantitative and (2) its restriction to MED1 and FUS condensates, and further developments are needed to provide more accurate and generally applicable predictions. Further, our simulations represent a single point of a multi-dimensional phase diagram, and broader sampling across different compositions of scaffold and clients will help disentangle the molecular interactions that determine mixing. In a future direction, sequence-based predictors could also be extended to account for compositional control of condensate partitioning ability arising from multiple scaffold components.^23,48,53,69^

## 4 Methods

### 4.1 Coarse-grained model potential

Proteins were modelled using a one-bead-per-amino acid residue representation in CALVA-DOS 2,^46^ which defines the potential as

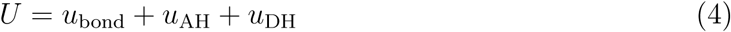

where *u*_bond_, *u*_AH_, *u*_DH_ are the bonded potential, AH potential and DH potential, respectively. RNA was modelled using a two-bead-per-nucleotide representation in CALVADOS-RNA,^48^ in which the potential is defined as

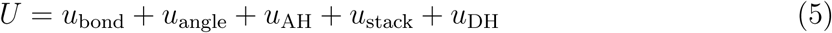

where *u*_angle_ is the angle potential, and *u*_stack_ is the stacking potential.

The bonded interaction between neighbouring beads is modelled by a harmonic potential,

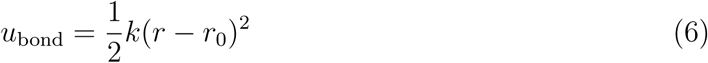

where *k* is the spring constant of 8033 kJ mol*^−^*^1^ nm*^−^*^2^ and *r*_0_ is the equilibrium bond length of 0.38 nm for proteins. In the bonded potential of RNA, the parameters for the backbone– backbone bonds are *k* = 1400 kJ mol*^−^*^1^ nm*^−^*^2^ and *r*_0_ = 0.59 nm, while those of backbone– base bonds are *k* = 2020 kJ mol*^−^*^1^ nm*^−^*^2^ and *r*_0_ = 0.54 nm. The angle potential is introduced only for the RNA backbones and is represented by

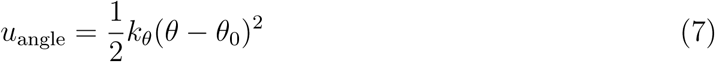

where *k_θ_* is the spring constant of 4.2 kJ mol*^−^*^1^ rad*^−^*^2^ and *θ*_0_ is the equilibrium angle of 3.14 rad. The non-bonded interactions are captured by stickiness-dependent interaction and electrostatic interaction. The stickiness-dependent interaction is represented by AH potential,^70^

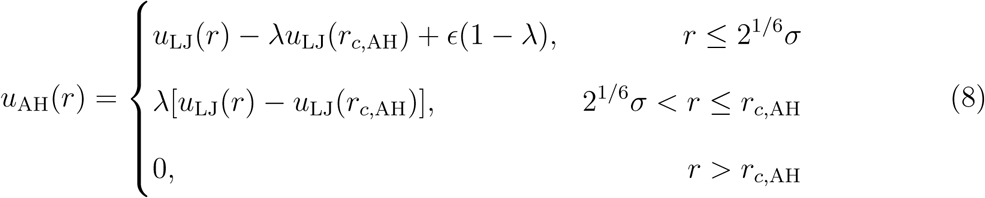

where *σ* = (*σ_i_* + *σ_j_*)*/*2, *λ* = (*λ_i_* + *λ_j_*)*/*2 for residues *i* and *j*, and the Lennard-Jones potential,

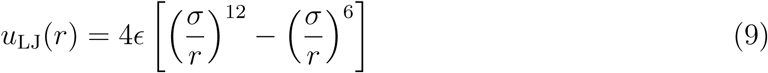

with *ɛ* = 0.8368 kJ mol*^−^*^1^, and *σ* obtained from Kim-Hummer model.^71^ The cutoff distance, *r_c,_*_AH_, was 2 nm. For RNA neighbouring bases, the stacking potential is represented as

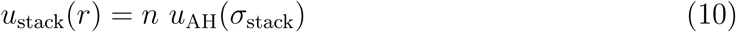

where *n* is the constant (*n* = 15) and *u*_AH_ is the same as AH potential but the volume parameter is *σ*_stack_ = 0.4 nm. The electrostatics is modelled by the DH potential,

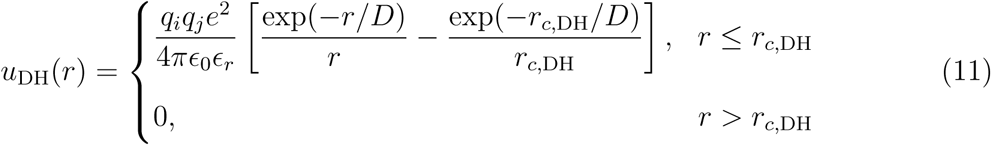

where *q* is the charge (±1), *ɛ*_0_ is vacuum permittivity, *ɛ_r_* is the dielectric constant of water, described by an empirical relationship as a function of the temperature *T*,^72^

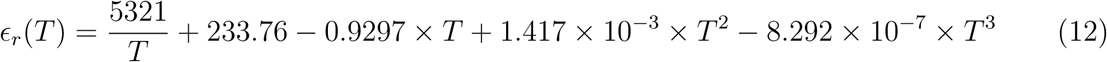

Debye length is 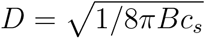 for Bjerrum length *B*(*ɛ_r_*), and ionic strength of monovalent ions *c_s_*. *r_c,_*_DH_ is the cutoff distance for DH potential and determined to be 4 nm.

### 4.2 Coarse-grained molecular dynamics simulation

IDR sequences were taken from the work by De La Cruz *et al.*^36^ and are shown in Fig S5. The number of scaffolds IDR chains in the simulation box was either 200 for MED1 or 586 for FUS, such that the number of total beads of these scaffolds IDR was approximately equal (∼120,500 beads). The RNA sequence was a generic homogeneous 250 bp RNA (polyRNA250), modelled as an unstructured single-stranded RNA.^48^ Each simulation system contained 34 polyRNA250 chains, which approximately neutralized the net charge of MED1 chains. The number of client IDR chains was 32 for all the chains. In simulations without RNA or without clients, RNA chains and/or client IDRs were removed while keeping all other components unchanged. The slab geometry of simulation box was 30 nm × 30 nm × 500 nm. In our simulations, the component concentrations were fixed by the simulation setup, but these settings did not necessarily quantitatively match the previously reported experimental conditions,^36^ nor did they lie on the same tie line in the three-component phase diagram space.

Each molecule was first constructed in an Archimedean-spiral conformation and then randomly placed within the simulation box to form the initial configuration. Subsequently, for fast simulation convergence, biased simulations were performed, with dragging external forces applied to proteins toward the centre of the box in the *z*-axis. Subsequently, unbiased simulations were performed for 16 *µ*s, with trajectories output every 10 ns, and the first 1 *µs* of simulation was discarded as equilibration. All simulations were performed at a temperature of 293 K, an ionic strength of 0.15 M, and histidine protonation state was set at pH 7.4. We used a Langevin integrator with a time step of 10 fs and a friction coefficient of 0.01 ps*^−^*^1^. All the simulations were performed using OpenMM 8.1.1,^73^ and simulation input files were generated using CALVADOS package.^74^

### 4.3 Concentrations of dilute and dense phases and partitioning free energy

In the analysis of slab-box simulations, we removed the drift of the condensate by computing its centre based on the density profile of the host IDR and repositioning it at the centre of the simulation box. Subsequently, density profiles for each component were computed along the longest box axis using a bin width of 1 nm. We determined the positions of the dense, dilute, and interfacial regions by fitting the semi-profiles of the scaffold IDR at *z <* 0 and 0 *< z* to the function of the interfacial density curve,^46,48,75^

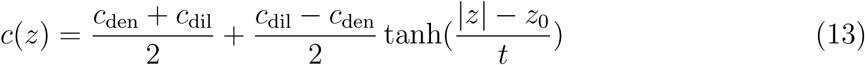

where *c*_den_, *c*_dil_, *t*, *z*_0_ are fitting parameters. The dense phase region was defined as |*z*| *< z*_0_ − *β*_den_*t* and the dilute phase as |*z*| *> z*_0_ + *β*_dil_*t*, with *β*_den_ = 3–4 and *β*_dil_ = 10. For each time frame, we averaged the densities of the two phases over the defined region to obtain the average concentrations of the phases. We computed the standard errors of the mean of the time series using a time-blocking approach (https://github.com/fpesceKU/BLOCKING).

Given the concentrations of the two phases, *c*_den_ and *c*_dil_, the partitioning free energy was defined as,

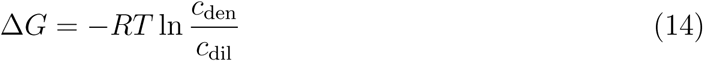

Due to finite size effects, concentrations below a few *µ*M are not reliably accessible in our simulations. Considering that *c*_dil_ = 1 *µ*M and *c*_den_ = 10^4^ *µ*M yield Δ*G* ∼ −6.9*RT*, we assigned |Δ*G*| = 8 *RT* to systems exhibiting substantial partitioning or exclusion. However, these values should be regarded as qualitative estimates rather than precise measurements.

### 4.4 Interaction energy within condensates

Provided that the condensate was positioned at the centre of the simulation box, we represented molecular interactions within the condensate by those of the central chain in the simulation box.^45^ At each time frame, we computed the centre of mass of the chains of interest, and selected the chain closest to the system centre as the central chain. Intermolecular interactions between this central chain and the surrounding beads were calculated using the residue-level potential functions (Eqs. 8 and 9). The central chain was re-identified at each frame, and the interaction energies were averaged over time. The standard error of the mean was estimated using the time-blocking approach.

### 4.5 Sequence-based predictor of client partitioning

Sequence-based predictor of client partitioning performs (1) assigning residue types, (2) generating sticker blocks from the sequence, (3) estimating block–block interaction, and (4) summing block interactions. In the first step, we read the sequence from the N-terminus and used the CALVADOS parameters to compute the average stickiness and charge over 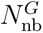 neighbouring residues (e.g. if 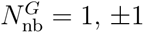 and 0 neighbouring residues are used). Stickiness was represented by the mean local aromatic parameter,

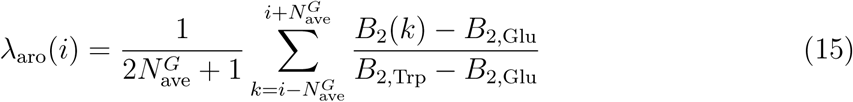

where *B*_2_(*i*), *B*_2,GLU_, *B*_2,TRP_ are the second virial coefficient of amino acid *i* without electrostatic potential, Glu and Trp, respectively. In CALVADOS 2, the stickiness parameter of Glu and Trp are the lowest and highest, respectively. Charge was represented by local charge parameter,

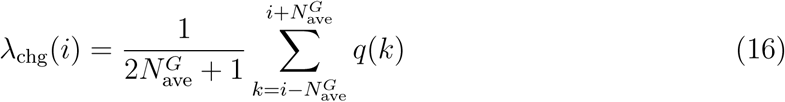

These averaged values were then reassigned to residue *i*, whereas the termini were excluded from this process. Next, residues were classified as aromatic, acidic, basic, or otherwise as spacer, using thresholds of stickiness, 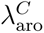, and charge, 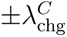, respectively. In the second step, we read the sequence from the N-terminus and grouped consecutive residues of the same sticker type into blocks. A block was initiated with two adjacent residues (*i*, *i* + 1) of the same type, and was extended by including additional neighbouring residues (*i* + 2, *i* + 3, …) of the same type. If a residue of a different type was encountered, or if the number of residues in a block exceeded the predefined maximum, 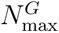, the current block was closed and a new block was initiated. In the third step, we computed the block–block interactions using Eq. 2.

We scanned the parameters related to sequence blocking and the energy scaling factor *α* so that *B*^tot^ qualitatively reproduces the trend of MED1/FUS partitioning. This scan led to the use of the following parameter set: 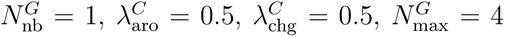, and *α* = 3.

## Data and code availability

The code and data for this paper are available at https://github.com/KULL-Centre/_2025_yasuda_partitioning. The code to run CALVADOS is available at https://github.com/KULL-Centre/CALVADOS.

## Acknowledgement

I.Y. acknowledges support from JSPS (KAKENHI, Grant Number JP23KJ1918, Japan), Marubun foundation (Exchange Research Grant, Japan) and JST (ACT-X, Grant Number JPMJAX24LJ, Japan). E.Y. acknowledges support from JST PRESTO (Grant JPMJPR22EE). The research was also supported by the PRISM (Protein Interactions and Stability in Medicine and Genomics) centre funded by the Novo Nordisk Foundation (NNF18OC0033950, to K.L.-L.).

## 5 Supporting Information

### 6 Prediction of intermolecular interactions using FINCHES

Pairwise interaction strength parameters (*ɛ* values) were obtained with FINCHES (version 0.1.2, beta public release). Calculations were performed under physiological conditions (293 K, pH 7.4, 0.150 M salt concentration), with both aliphatic weighting and charge weighting enabled. *ɛ* values were computed between client proteins and FUS, as well as between client proteins and MED1, considering only the intrinsically disordered regions of the proteins.

**Table S1:** Thresholds used in client classification using sequence-based predictors, (*α*_MED1_ and *α*_FUS_), and the corresponding Matthews correlation coefficients (MCCs). The MCCs were computed for four-class classification (common client, MED1 client, FUS client, and non-client).

| | $\alpha_{\text{MED1}}$ | $\alpha_{\text{FUS}}$ | MCC |
| --- | --- | --- | --- |
| Block | $-4 \times 10^9$ | $-2 \times 10^5$ | 0.18 |
| Finches | 110 | 2 | 0.15 |

**Figure S1:**
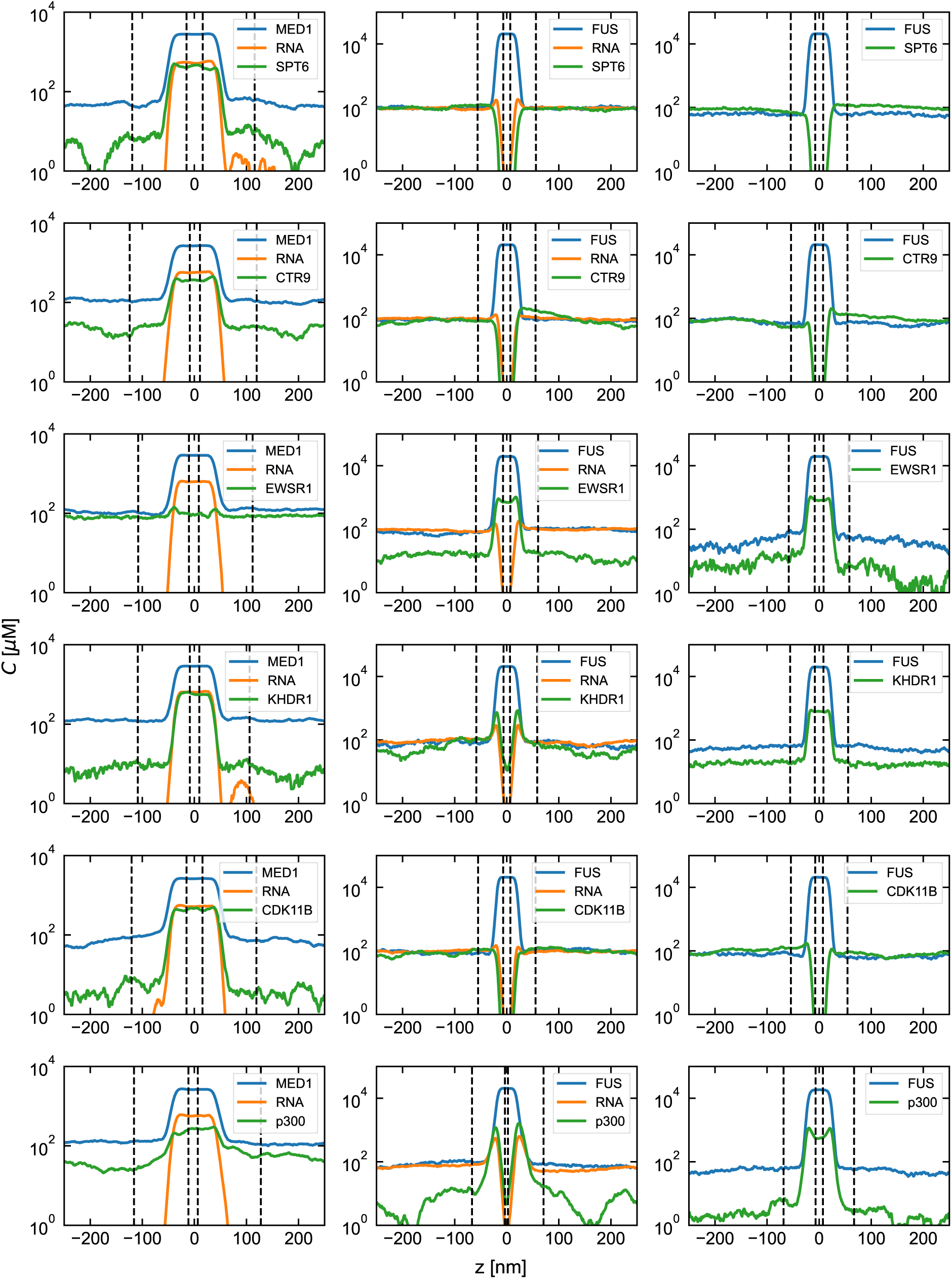
Density profiles showing component density, *C*, along the longest box axis, *z*. Dashed lines indicate the boundaries of the dense and dilute phases.

**Figure S2:**
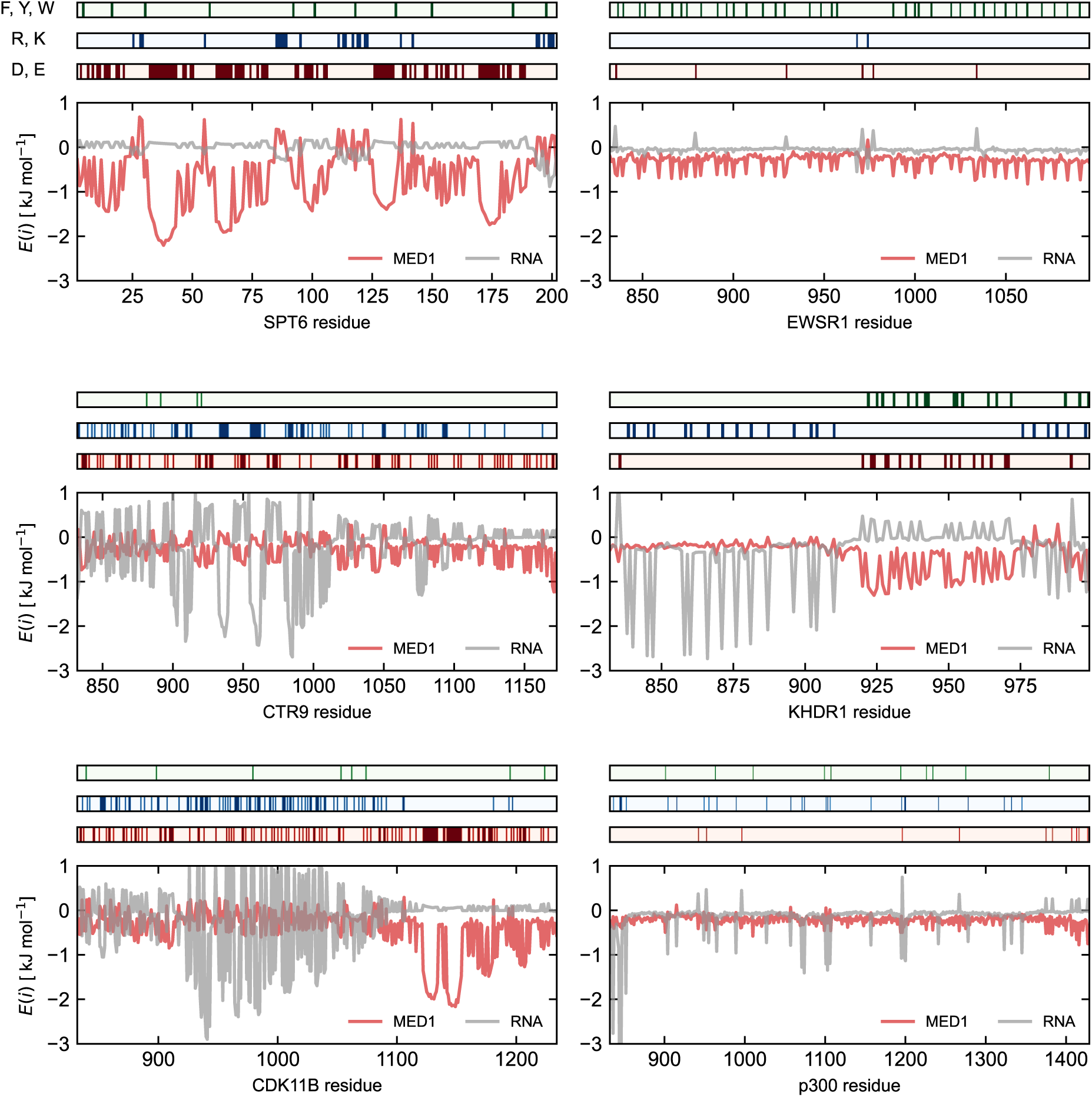
Analysis of client interaction energy per residue within MED1 condensates.

**Figure S3:**
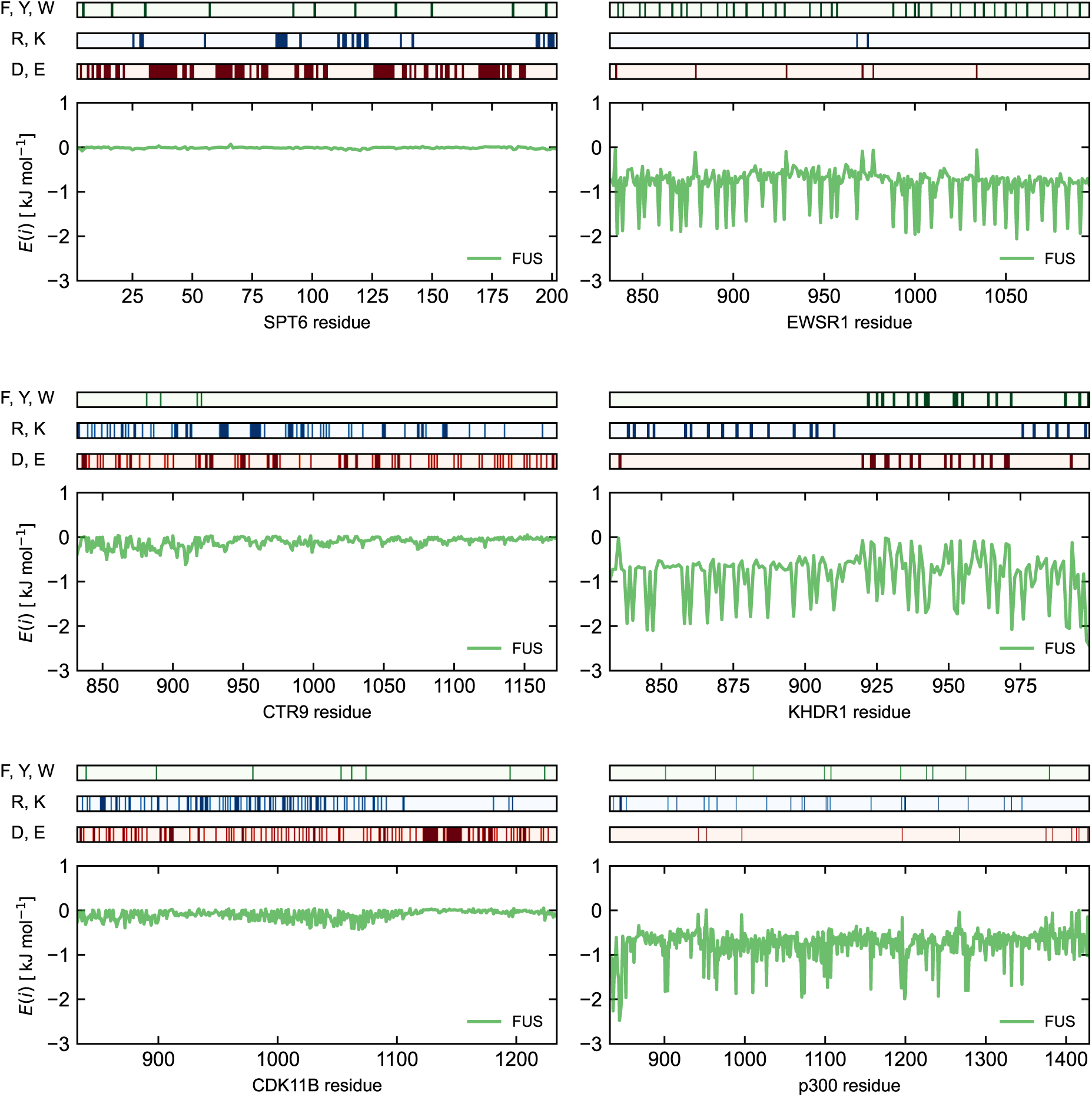
Analysis of client interaction energy per residue within FUS condensates.

**Figure S4:**
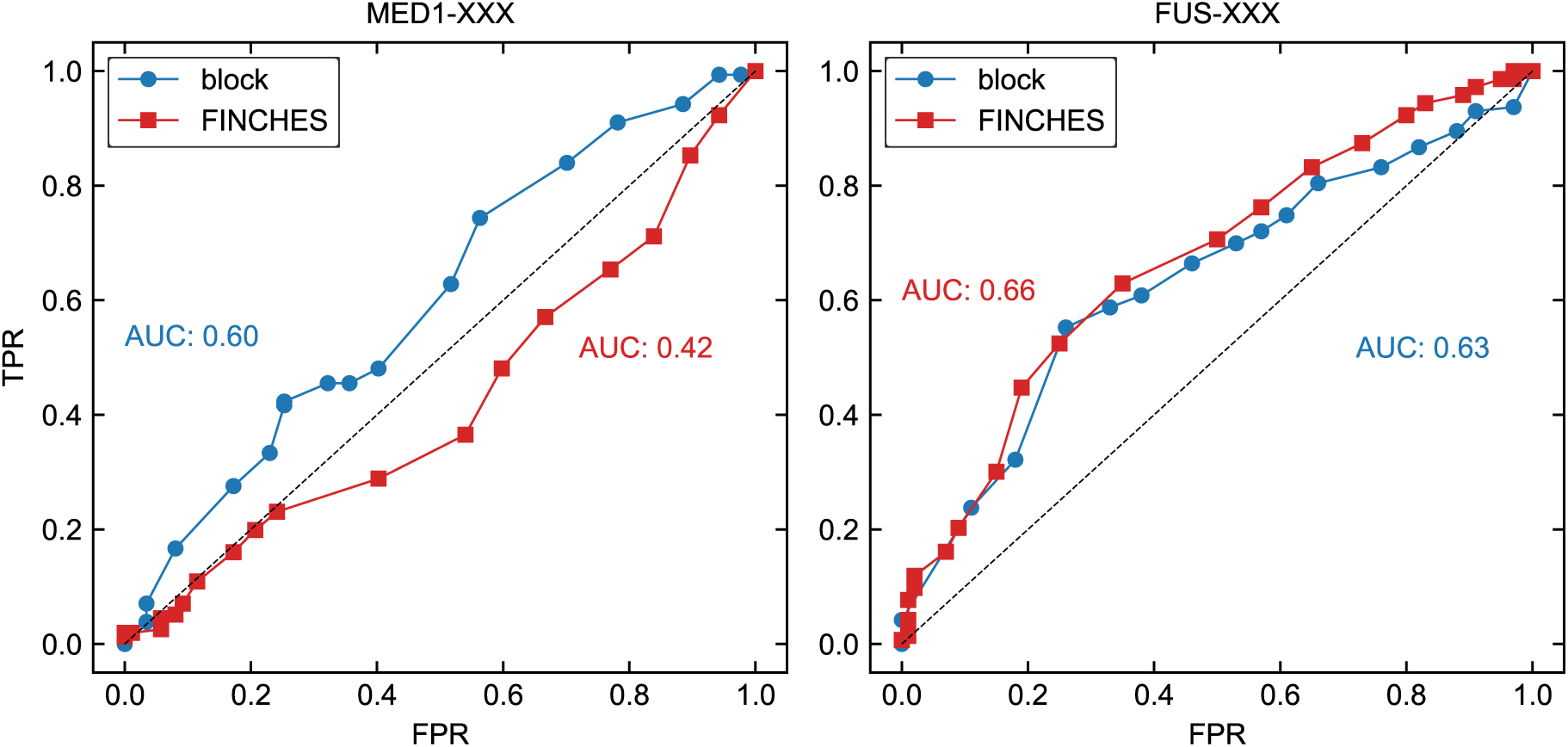
Receiver operating characteristic (ROC) curves for our blocking predictor (Block) and FINCHES in classifying client proteins. ‘MED1–XXX’ denotes MED1–client interactions, while ‘FUS–XXX’ denotes FUS–client interactions. For MED1 clients, the ‘common’ and ‘MED1’ groups were treated as true positives, whereas the ‘FUS’ group was treated as false. For FUS clients, the ‘FUS’ and ‘common’ groups were treated as true positives, while the ‘MED1’ group was treated as false. True positive rate (TPR) and false positive rate (FPR) were calculated by varying the threshold of the interaction score. The area under the curve (AUC) was computed numerically using the trapezoidal rule.

**Figure S5:**
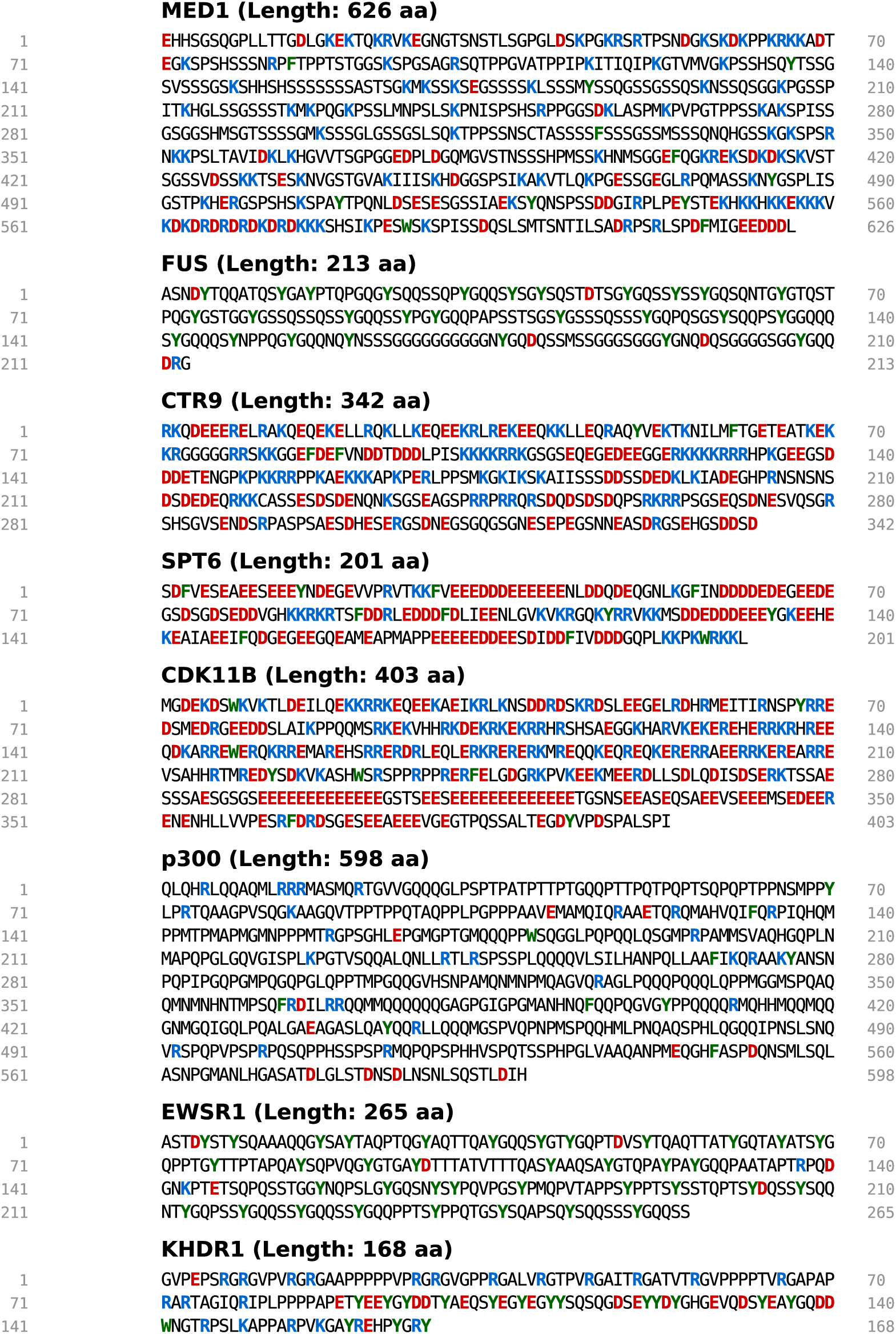
Protein sequences used in MD simulations. Coloured residues indicate acidic residues (red), basic residues (blue), and aromatic residues (green).

